# High-throughput Identification of Eukaryotic Parasites and Arboviruses in Mosquitoes

**DOI:** 10.1101/2021.01.12.426319

**Authors:** Matthew V. Cannon, Haikel N. Bogale, Devika Bhalerao, Kalil Keita, Denka Camara, Yaya Barry, Moussa Keita, Drissa Coulibaly, Abdoulaye K. Kone, Ogobara K. Doumbo, Mahamadou A. Thera, Christopher V. Plowe, Mark A. Travassos, Seth R. Irish, Joshua Yeroshefsky, Jeannine Dorothy, Brian Prendergast, Brandyce St. Laurent, Megan L. Fritz, David Serre

**Author notes:** correspondence to, University of Maryland School of Medicine, Institute for Genome Sciences, Department of Microbiology & Immunology, HSFIII, 670 West Baltimore Street, Room 3111, Baltimore, MD 21201. Deceased (Ogobara K. Doumbo). **Author contributions** MVC, BSTL, MF and DS designed the study. BSTL, MF, KK, DC, YB, MK, DC, AKK, OKD, MAT, CVP, MT, SI, JD and BP collected and/or provided access to the mosquito samples. MVC, HB, BSTL and DB performed the laboratory experiments. MVC analyzed the data. MVC and DS prepared the manuscript. All authors read and approved the final manuscript.

## Abstract

Vector-borne pathogens cause many human infectious diseases and are responsible for high mortality and morbidity throughout the world. They can also cause livestock epidemics with dramatic social and economic consequences. Due to the high costs, vector-borne disease surveillance is often limited to current threats, and the investigation of emerging pathogens typically occur after the reports of clinical cases. Here, we use high-throughput sequencing to detect and identify a wide range of parasites and viruses carried by mosquitoes from Cambodia, Guinea, Mali and Maryland. We apply this approach to individual *Anopheles* mosquitoes as well as pools of mosquitoes captured in traps; and compare the outcomes of this assay when applied to DNA or RNA. We identified known human and animal pathogens and mosquito parasites belonging to a wide range of taxa, insect Flaviviruses, and novel DNA sequences from previously uncharacterized organisms. Our results also revealed that analysis of the content of an entire trap is an efficient approach to monitor and identify potential vector-borne pathogens in large surveillance studies, and that analyses of RNA extracted from mosquitoes is preferable, when possible, over DNA-based analyses. Overall, we describe a flexible and easy-to-customize assay that can provide important information for vector-borne disease surveillance and research studies to efficiently complement current approaches.

## Introduction

Different arthropods can, during a blood feeding, transmit viruses, protists and helminths to humans (1). These organisms cause some of the most prevalent human infectious diseases, including malaria, dengue, schistosomiasis or Chagas disease, and are responsible for more than 700,000 human deaths worldwide every year (2–4). Vector-borne diseases are also responsible for some of the most alarming recent epidemics in the western hemisphere, either due to the emergence of new pathogens (e.g., Zika (5, 6)), the reemergence of historically important pathogens (e.g., Yellow Fever (7)) or the expansion of diseases beyond their historical ranges (e.g., West Nile (8) and Chikungunya (7)). In addition to this burden on human health, many vector-borne diseases affect domesticated animals (e.g., heartworms (9, 10)), livestock (e.g., Theileriosis (11, 12)) and wild animals (e.g., avian malaria (13, 14)). Some of these animal diseases have dramatic economic consequences in endemic areas (11, 12), while others are zoonotic diseases, further affecting human populations (15–20).

Efficient vector-borne disease surveillance is critical for reducing disease transmission and preventing outbreaks. Past elimination campaigns for vector-borne diseases, usually targeting a specific human pathogen, have often relied on entomological approaches such as widespread insecticide spraying and disruption of larval habitats (21, 22). To be successful, such efforts need to be guided by detailed knowledge of the parasites’ and vectors’ distributions. Unfortunately, current entomological surveillance approaches are extremely resource-intensive: the collection of samples is time consuming requiring trained personnel, vector species identification is laborious, and the detection of pathogens is expensive since hundreds of mosquitoes typically need to be screened to identify a few infected ones. Consequently, public health officials and vector biologists typically focus on monitoring only a few specific pathogens associated with the most current threats. These constraints are particularly problematic as they hamper the early detection of emerging pathogens and vector surveillance is often implemented in response to reports of clinical cases rather than preventively.

We have recently described a sequencing-based method using high-throughput amplicon sequencing to detect known and previously uncharacterized eukaryotic parasites from biological samples in a comprehensive, high-throughput and cost-efficient manner (23). Here, we present the application of this approach to characterize eukaryotic parasites and arboviruses from more than 900 individual *Anopheles* mosquitoes collected in Cambodia, Guinea and Mali, as well as from 25 pools of mosquitoes captured in CDC CO_2_-baited light traps in Maryland, USA. We also compare the performance of the assay when screening DNA and RNA from the same samples. Overall, our study demonstrates how this sequencing-based assay could significantly improve monitoring of human and animal vector-borne pathogens.

## Methods

### Samples

We analyzed a total of 930 individual mosquitoes, as well as 25 pools each containing 50-291 mosquitoes (2,589 total) (**Table 1** and **Supplemental Tables 1-3**).

**Table 1.**
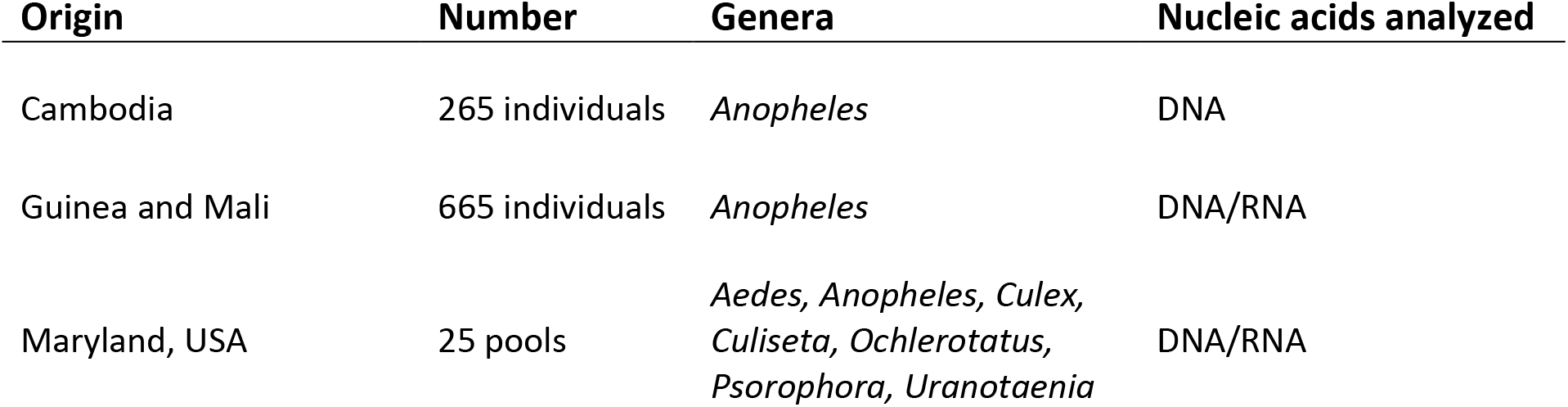
Sample summary.

| Origin | Number | Genera | Nucleic acids analyzed |
| --- | --- | --- | --- |
| Cambodia | 265 individuals | *Anopheles* | DNA |
| Guinea and Mali | 665 individuals | *Anopheles* | DNA/RNA |
| Maryland, USA | 25 pools | *Aedes, Anopheles, Culex, Culiseta, Ochlerotatus, Psorophora, Uranotaenia* | DNA/RNA |

First, we analyzed DNA previously extracted from 265 individual *Anopheles* mosquitoes collected in the Cambodian provinces of Pursat, Preah Vihear, and Ratanakiri (24). These mosquitoes were collected using cow- or human-baited tents, human landing collections, CDC light traps and barrier-screen fences and immediately preserved by desiccation upon collection. These 265 *Anopheles* mosquitoes represent 22 different species collected between July and August of 2013 (see **Supplemental Table 1** for details).

Second, we included DNA samples from 81 individual mosquitoes collected in Bandiagara, Mali. DNA from these samples was extracted using Chelex^®^ 100 (Bio-Rad) after incubation of bisected and homogenized mosquitoes in 1% saponin in PBS.

Third, we extracted DNA from 584 individual *Anopheles* mosquitoes collected in six sites in Guinea and preserved in ethanol immediately upon collection. These mosquitoes were collected by human landing catch and pyrethrum spray (**Supplemental Table 2**). Each mosquito was homogenized in 200 μl ATL/proteinase K solution using five RNase-free 1 mm zirconium oxide beads in a TissueLyser II for 12 minutes at 20 m/s. We centrifuged the solution at 2500 rpm for three minutes and incubated them at 55°C for one hour. We performed a second homogenization step for four minutes at 20 m/s followed by a final incubation at 55°C overnight. We then isolated DNA using the Qiagen DNeasy 96 Blood & Tissue Kit according to the manufacturer’s instruction and eluted DNA from each sample in 200 μl.

Finally, we analyzed 25 pools of mosquitoes collected throughout Prince George’s county (Maryland, USA) by the Maryland Department of Agriculture using CO_2_-baited light traps (**Supplemental Table 3**). Each pool contains all mosquitoes from one light-trap (^~^50-291 mosquitoes) and was stored at room temperature for up to 24 hours before long-term storage at −20°C. We homogenized each pool of mosquitoes using a Qiagen TissueLyser II with Teenprep Matrix D 15 ml homogenization tubes (MP Biomedicals) and isolated successively RNA and DNA from each sample using the RNeasy PowerSoil Total RNA kit (Qiagen) with the RNeasy PowerSoil DNA Elution Kit and a final elution volume of 100 μl.

### Evaluation of Arbovirus primers

We tested universal flavivirus primers retrieved from the literature (25) on West Nile (n=3), Zika (n=2) and Dengue (n=2) viral RNAs obtained from the American Type Culture Collection (ATCC). We synthesized cDNA from 2 μL of RNA using M-MLV reverse transcriptase (Promega) and random hexamers, and amplified the resulting cDNA with GoTaq^®^ DNA polymerase (Promega) under the following conditions: initial two-minute denaturing step at 95°C followed by 40 cycles of 95°C for 30 seconds, 50°C for 30 seconds and 72°C for 40 seconds. A final extension at 72°C for ten minutes was followed by incubation at 4°C. We ran the products on an agarose gel to determine whether each virus RNA was amplifiable.

### PCR amplification of pathogen nucleic acids before high-throughput sequencing

First, we synthesized cDNA using M-MLV reverse transcriptase (Promega) and random hexamers from either 1) 2 μl of the nucleic acids isolated from the Guinean mosquitoes (i.e., using RNA carried over during the DNA extraction), or 2) 3 μL of RNA extracted from the pools of Maryland mosquitoes.

Then, we amplified DNA and cDNA (when available) from each sample, as well as from 176 no-DNA controls, with a total of 11 primer pairs, each targeting a specific taxon known to contain human pathogens (**Table 2**). For each primer pair, we amplified DNA and cDNA using GoTaq^®^ DNA polymerase (Promega) under the following conditions: initial denaturing step at 95°C followed by 40 cycles of 95°C for 30 seconds, 50°C for 30 seconds and 72°C for 30 seconds. A final extension at 72°C for ten minutes was followed by incubation at 4°C. All primers used in these taxon-specific PCRs included 5’-end tails to serve as priming sites for a second PCR. We then pooled all PCR products generated from one sample and performed a second PCR using primers targeting these tails to incorporate, at the end of each amplified molecule, i) a unique oligonucleotide “barcode” specific to each sample and ii) DNA sequences complementary to the Illumina sequencing primers (26, 27) (**Supplemental Figure 1**). Finally, we pooled together the resulting barcoded libraries and sequenced them on an Illumina sequencer to generate an average of 12,703 paired-end reads of 251 or 301 bp per sample.

**Table 2.**
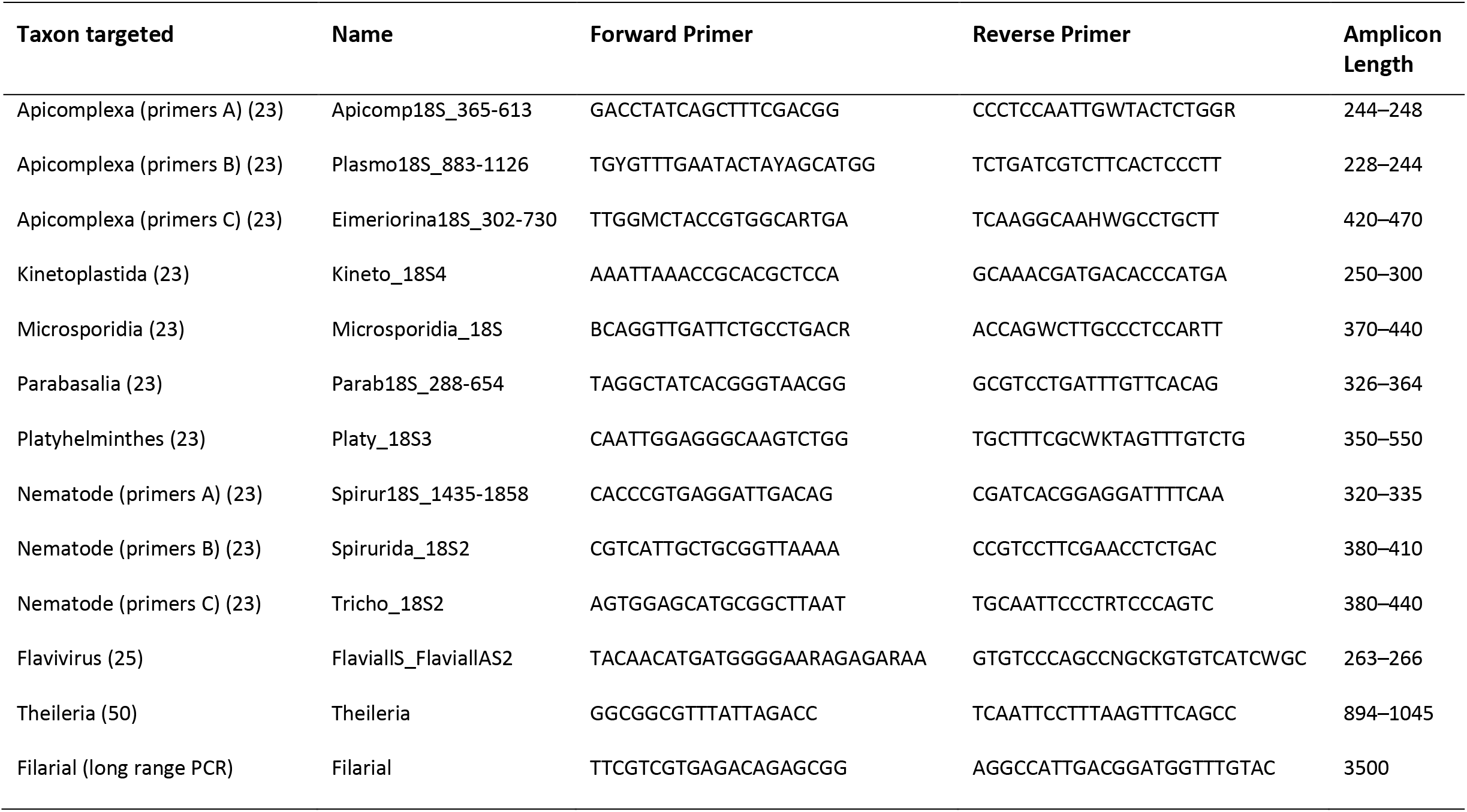
Primers used.

### Bioinformatic analyses

We first separated the reads generated from each sample according to their unique barcodes and merged the overlapping ends of each read pair using PANDAseq (28) to generate consensus DNA sequences and correct sequencing errors (that disproportionally occur at the end of the reads). Note that all primers were designed to amplify DNA sequences shorter than ^~^450 bp, allowing overlap of at least 50 bp between paired-end reads. All read pairs that did not merge correctly were discarded from further analyses. We identified and trimmed the primer sequences from each read and eliminated all consensus sequences shorter than 100 bp as they likely represent experimental artefacts (e.g., PCR chimeras and primer dimers). Using reads from all samples together, we recorded how many unique DNA sequences were obtained and how many reads carried each of these unique DNA sequences. Sequences observed less than ten times in the entire dataset were omitted as they likely resulted from PCR or sequencing errors (26). We then compared each unique DNA sequence to all sequences deposited in the NCBI nt database using BLAST (29) and used custom pipelines (https://github.com/MVesuviusC/2020MosquitoSurveillancePaper) to retrieve the taxonomic information associated with the most similar sequence(s). For each sample, only sequences with at least 10 reads and more than 70% identity with an annotated NCBI sequence over the entire sequence length were further considered. This identity cutoff, while low, allows inclusion of results from highly genetically divergent organisms which can then be scrutinized further. This is critical when identifying species without closely related sequences available. If DNA sequences from multiple species were equally similar to one of our sequences, we recorded all corresponding species names. Finally, we summarized, for each mosquito, the parasite species or virus identified, the percentage identity between the reads and the most similar NCBI sequence(s), and the number of reads supporting the identification in this sample.

### Phylogenetic analyses

To better characterize specific DNA sequences with ambiguous species identification, we analyzed these sequences together with orthologous sequences from closely related species. Briefly, we used PrimerTree (26) to retrieve NCBI orthologous DNA sequences from all species of the targeted taxon. We aligned these sequences with the DNA sequence(s) amplified from the mosquito(es) using MAFFT (30) and reconstructed neighbor-joining trees using MEGA (31) to estimate the phylogenetic position of the amplified DNA sequences.

### Further determination of taxonomical assignments

To improve species identification when multiple species had identical DNA sequences, or improve phylogenetic analyses of unknown sequences, we amplified and sequenced specifically chosen DNA loci from pathogens within the same mosquitoes.

For differentiating *Theileria* species, we used previously published primers (GGCGGCGTTTATTAGACC, TCAATTCCTTTAAGTTTCAGCC) to amplify an informative portion of the 18S rRNA gene [31] using DNA from 19 samples identified as *Theileria* positive by high-throughput sequencing. Amplification was conducted under the following conditions: initial denaturing step at 95°C for two minutes followed by 40 cycles of 95°C for 30 seconds, 50°C for 30 seconds and 72°C for 40 seconds. A final extension at 72°C for five minutes was followed by incubation at 4°C. Since gel electrophoresis revealed off-target amplification (multiple bands), we used a Pasteur pipette to collect a core from the agarose gel, corresponding to the expected 900 bp PCR product, and dissolved it in 100 μl of water at 60°C for 20 minutes. We then re-amplified 10 μl of this DNA using 35 PCR cycles with the same conditions. After gel electrophoresis, we treated the PCR reaction with 0.046 μl of Exonuclease I (NEB) and 0.4625 μl of Shrimp alkaline phosphatase (Affymetrix) at 37°C for 30 minutes, with a final five-minute inactivation step at 95°C. We then Sanger sequenced each PCR product in both directions using the forward and reverse primers. We manually trimmed the reads and merged them using Flash (32). We aligned the reads, along with known *Theileria* sequences from the NCBI nucleotide database, using MAFFT (30, 33) and generated a neighbor joining tree with 500 bootstraps and plotted it in MEGA7 (34).

To identify the species of the filarial worms detected in two individual mosquitoes, we designed primers to amplify a 3.5 kb portion of the mitochondrial DNA. Briefly, we downloaded all available filarial worm (Filarioidea) mitochondrial sequences from the NCBI nucleotide database, aligned them, generated a consensus sequence and designed primers using primer3 (35). We then used these primers (TTCGTCGTGAGACAGAGCGG, AGGCCATTGACGGATGGTTTGTAC) to amplify DNA from the two positive mosquitoes using the Expand™ Long Range dNTPack kit (Sigma) using the following conditions: initial denaturing step at 95°C for two minutes followed by 45 cycles of 92°C for 30 seconds, 55°C for 30 seconds and 68°C for five minutes. A final extension at 68°C for ten minutes was followed by incubation at 4°C. We then performed a second PCR to add 10 bp barcodes to the 5’ end of both forward and reverse primers to allow differentiating both samples after sequencing. The two barcodes differed by 8 and 7 bases for the forward and reverse primers, respectively, with no more than 2 identical bases in a row (**Supplemental Table 4**). For this second PCR, we used the following conditions: initial denaturing step at 95°C for two minutes followed by 10 cycles of 92°C for 30 seconds, 55°C for 30 seconds and 68°C for five minutes. A final extension at 68°C for ten minutes was followed by incubation at 4°C. We purified the amplicons using AMPure XP beads (Beckman Coulter) (2:1 DNA:beads ratio) and then combined equimolar amounts of each barcoded PCR product before circular consensus sequencing on a PacBio Sequel. We then generated a consensus sequence for each sample and aligned these sequences to known nematode mitochondrial sequences using Mafft (30) and generated a neighbor joining tree in MEGA (36).

### Assessment of the dynamics of viral and mosquito RNA degradation

To assess the dynamics of viral RNA degradation over time, we analyzed colony *Culex pipiens* mosquitoes known to carry *Culex flavivirus*. The colony was initiated from diapausing adult *Culex pipiens* that were collected from Oak Lawn and Des Plaines, IL, on 2/8/10. These two collections were combined to make one colony, which was determined to be *Culex flavivirus* positive according reverse transcriptase PCR (25). We examined three pools of five mosquitoes for each condition (i.e., stored with no preservative, in ethanol or in RNAlater (Invitrogen)) and at each time point (i.e., fresh, after two-week or after four-week storage at room temperature). After 0, 2 or 4 weeks at room temperature, the mosquitoes were stored at −80°C until RNA isolation. We isolated RNA from each pool of mosquitoes using Qiazol (Qiagen) and eluted into 50 μl. We synthesized cDNA from 7 μl of RNA using m-MLV (Promega) with random hexamers for PCRs using *Culex* primers and, separately, on 2 μl of RNA for PCRs using flavivirus primers.

For each pool of 5 *Culex* mosquitoes from the *Culex flavivirus*-infected colony, we performed quantitative reverse transcriptase PCR (qRT-PCR) to quantify the amount of mosquito and viral RNAs using the primers Culex_flavivirus_3F (TGCGAARGATCTDGAAGGAG) - Culex_flavivirus_3R (CACGCACAACAAGACGATRA) targeting the virus sequence, and Culicinae_Cox1_379_F (AYCCHCCTCTTTCATCTGGA) - Culicidae_Cox1_670_R (CCTCCTCCAATTGGRTCAAAG) targeting mosquito RNA. We used Perfecta SYBR green PCR mastermix (Quantabio) with the following conditions: initial 15-minute denaturing step at 95°C followed by 40 cycles of 95°C for 30 seconds, 55°C (Culex primers) or 50°C (flavivirus primers) for 30 seconds and 72°C for one minute (Culex primers) or 40 seconds (flavivirus primers). We performed standard cycle threshold and melt curve analysis afterwards using default settings.

## Results

### Amplicon sequencing for high-throughput characterization of microorganisms in mosquitoes

We analyzed 265 *Anopheles* mosquitoes collected in Cambodia, 665 *Anopheles* mosquitoes collected in Guinea and Mali as well as the content of 25 light traps, each containing 50-291 mosquitoes, collected in Maryland, USA. We screened each sample for a wide range of eukaryotic parasites using 10 primer sets designed to amplify DNA from all species of the taxa known to include human pathogens: Apicomplexans, Kinetoplastids, Parabasalids, nematodes, Platyhelminthes and Microsporidians (**Table 2**). We also screened RNA extracted from the individual African *Anopheles* and from the pools of mosquitoes from Maryland for flaviviruses (see Materials and Methods, **Table 2**). After taxon-specific amplification, we pooled all PCR products generated from the same mosquito together, barcoded them and sequenced all libraries to generate an average of 12,703 paired-end reads per sample (**Supplemental Figure 1**). After merging read pairs, stringent quality filters and removal of the products of off-target amplification (e.g., *Anopheles* and bacteria DNA sequences), we obtained 61,177 unique DNA sequences, each represented by ten reads or more, and accounting in total for 6,796,105 reads (**Supplemental Table 5**). These sequences were amplified with all primers and from a total of 185 samples: 42 out of 265 Cambodian mosquitoes (16%), 120 out of 665 African mosquitoes (18%), and 23 out of the 25 pools (92%) of mosquitoes collected in Maryland were positive for at least one of the taxa tested. On average, each sequence was supported by 1,306 reads per sample (range: 10-43,440). By contrast, out of 176 negative controls, only 12 (7%) yielded any sequence from the targeted taxa and those were represented by 213 reads on average (range: 10-3,539).

### Identification of eukaryotic parasites

We retrieved DNA sequences identical to sequences previously amplified from *Theileria* parasites from 22 African and 15 Cambodian mosquitoes, as well as from seven of the Maryland traps. *Theileria* sequences were successfully amplified with both the Apicomplexa and Eimeronia primer pairs. All samples positive for *Theileria* with the Eimeronia primers were also positive with the Apicomplexa primers. On the other hand, the Eimeronia primers provided sufficient information to assign each sequence to a single species, while the sequences amplified with Apicomplexa primers were unable to differentiate among the *Theileria* species (see also below). We detected sequences identical to *Plasmodium falciparum* in eight African samples and two Cambodian samples, while sequences most similar (82.0%-99.5% identity) to bird *Plasmodium* species were amplified from 20 of the 25 traps in Maryland (**Table 3**). We also amplified a sequence that was identical to several *Babesia* species (100% identity) in one trap by two different primer pairs. Finally, we detected DNA from a known apicomplexan parasite of mosquitoes, *Ascogregarina barretti* (37), in two of the traps.

**Table 3.**
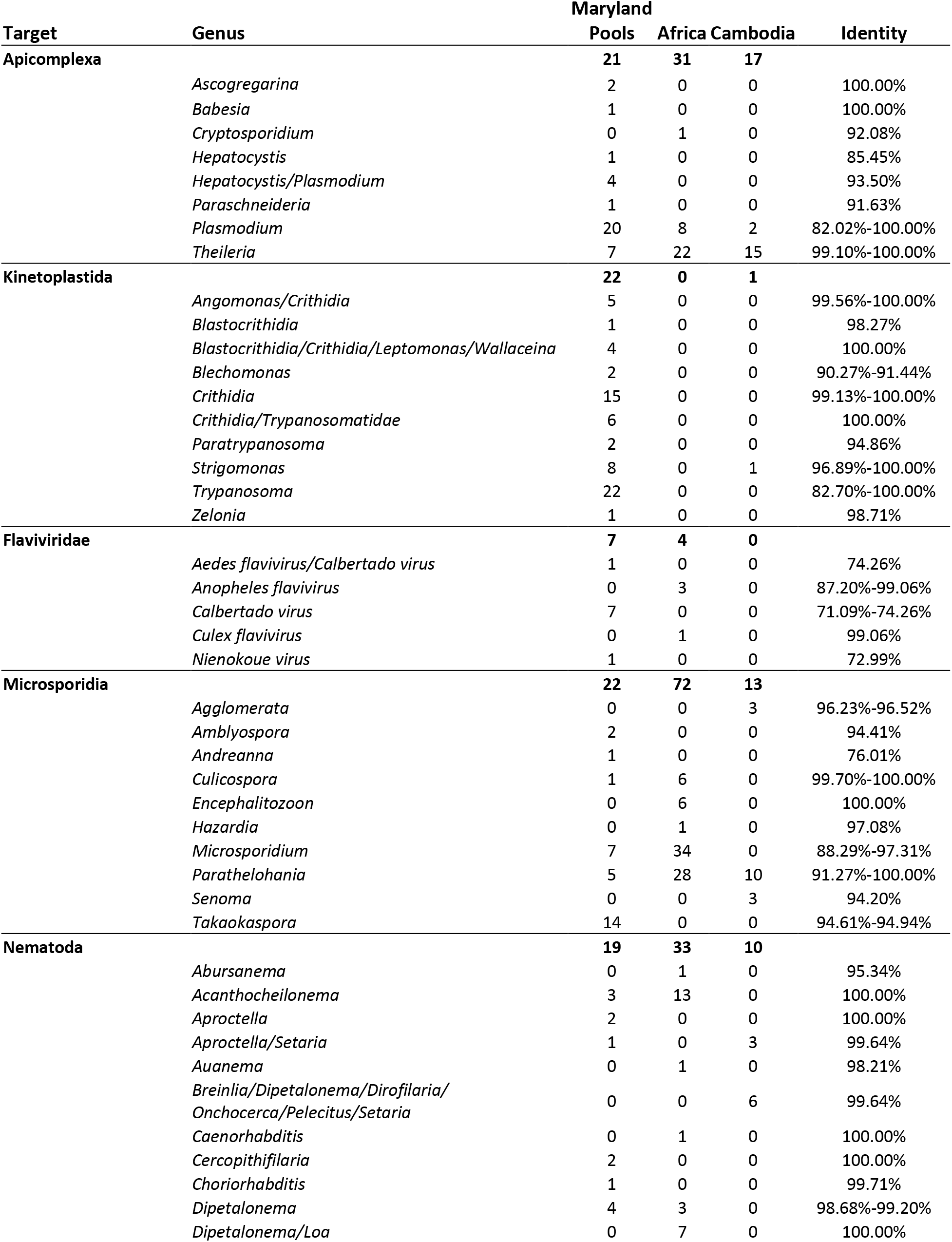
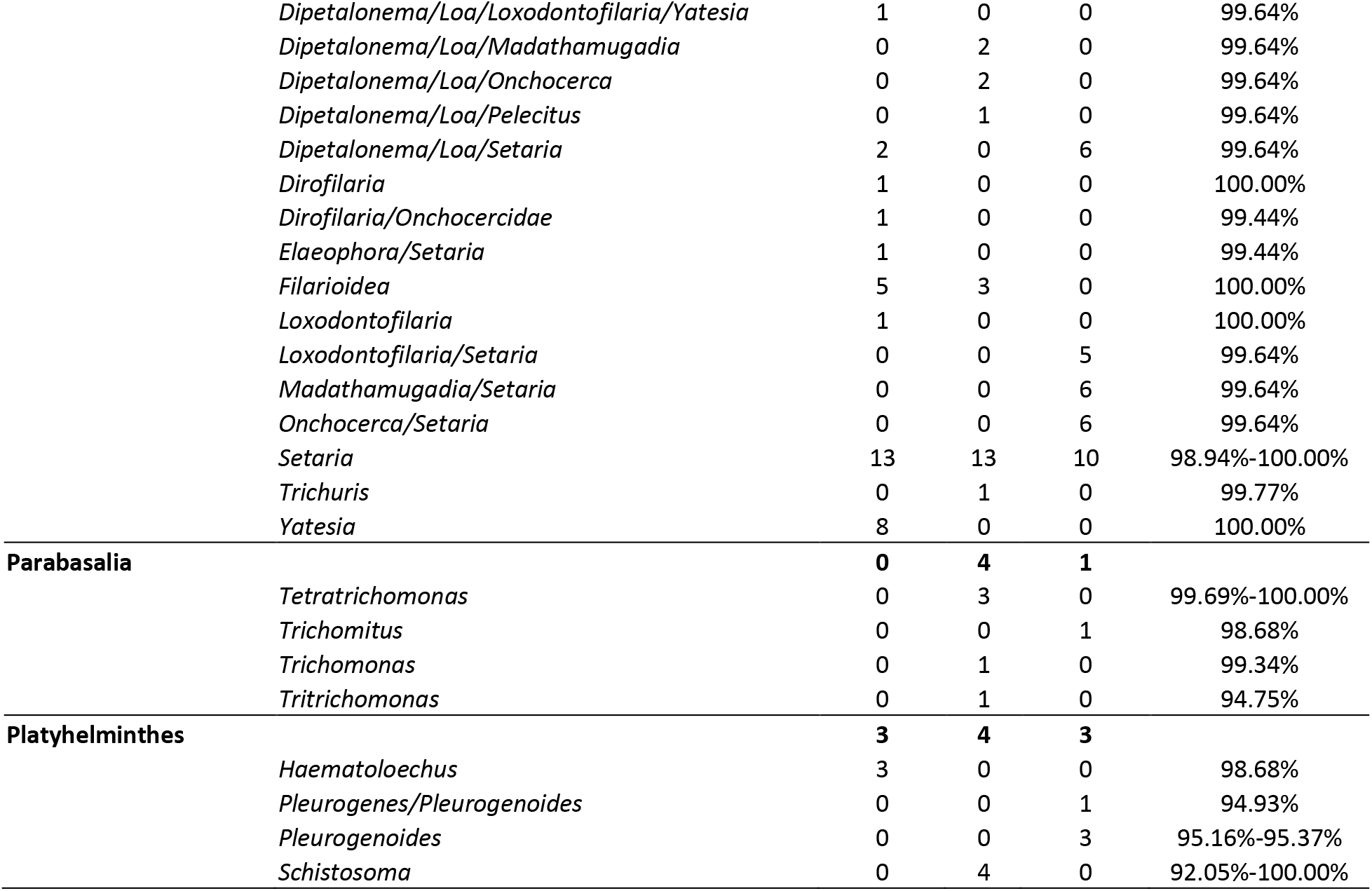
Genera amplified from each group of samples.

From all individual mosquitoes, only one Cambodian *Anopheles* yielded a Kinetoplast sequence that was most similar to *Strigomonas culicis* (96.9% identity). By contrast, 22 of the traps were positive for Kinetoplasts, yielding sequences similar to sequences from *Angomonas*, *Blastocrithidia, Blechomonas, Crithidia, Leptomonas, Paratrypanosoma, Strigomonas, Trypanosoma, Trypanosomatidae, Wallaceina* or *Zelonia* (with 90.3%-100% identity, except for one sequence that matched *Trypanosoma theileri* at 82.7% identity) (**Table 3**).

Many sequences were amplified using the Microsporidia primers: 72 African mosquitoes were positive with sequences similar or identical to *Culicospora*, *Encephalitozoon*, *Hazarida, Microsporidium* and *Parathelohania* (88.3%-100% identity), while 13 Cambodian samples yielded sequences similar or identical to *Agglomerata*, *Parathelohania* and *Senoma* (91.3%-100% identity) (**Table 3**). Twenty-two traps also yielded Microsporidia sequences closely matching those of *Amblyospora, Andreanna, Culicospora, Microsporidium, Parathelohania* and *Takaokaspora* (with 76.0%-100% identity).

Regarding parasites from the Parabasalia group, four African mosquitoes were positive for *Tetratrichomona, Trichomonas* or *Tritrichomonas* with high sequence similarity (94.8%-100%) while a single Cambodian mosquito was positive for *Trichomitus* (98.7% identity). No Parabasalia were detected in the Maryland traps.

We detected Platyhelminthes sequences in four African mosquitoes, all similar to *Schistosoma mansoni* (92.1%-100% identity). Three Cambodian mosquitoes yielded sequences most similar to those of either *Pleurogenoides or Pleurogenes* (94.9%-95.4% identity). Three traps in Maryland were positive for Platyhelminthes, with sequences most similar to *Haematoloechus* (98.7% identity).

The taxonomic resolution of the nematode primers was lower than that of the other taxon-specific primer pairs and the amplified sequences often matched multiple species (or even genera). We amplified nematode sequences from 33 African *Anopheles*, including sequences most similar to *Abursanema*, *Acanthocheilonema*, *Auanema*, *Caenorhabditis*, *Dipetalonema*, *Filarioidea*, *Loa*, *Loxodontofilaria*, *Madathamugadia*, *Onchocerca*, *Pelecitus*, *Setaria* or *Trichuris,*although the sequence similarity (95.3-100%) clearly indicated that, in some cases, the exact identity of the species was unknown (see also below). Ten Cambodian mosquitoes were positive for *Setaria digitata* (100% identity) while other mosquitoes yielded sequences that matched *Setaria* and one or more of the following genera: *Aproctella*, *Breinlia*, *Dipetalonema*, *Dirofilaria*, *Loa*, *Loxodontofilaria*, *Madathamugadia*, *Onchocerca*, *Pelecitus*. Nineteen different traps from Maryland produced nematode sequences with particularly high read counts of *Setaria*, *Yatesia* and *Dirofilaria* sequences (98.9% - 100% identity). Other genera detected in the traps included *Acanthocheilonema*, *Aproctella*, *Cercopithifilaria*, *Choriorhabditis*, *Dipetalonema*, *Elaeophora*, *Filarioidea*, *Loa*, *Loxodontofilaria*, *Onchocercidae*.

Overall, using this single assay, we screened over 3,500 mosquitoes from three geographic locations and identified DNA sequences from numerous microorganisms encompassing six classes, 12 orders and 23 families (**Table 3**).

### Identification of Flaviviruses in mosquitoes

To detect and identify flaviviruses, we used a primer pair predicted *in silico* to amplify a wide range of flaviviruses, including all known human pathogens (25), and we validated that these primers successfully amplified cDNA generated from West Nile, Zika and Dengue viruses. Out of 665 individual African mosquitoes, three were positive for viruses most similar to *Anopheles flavivirus variants 1* and *2* (87.2%-99.1% identity) (**Figure 1 and Supplemental Figure 2**) and one was positive for a virus similar to *Culex flavivirus* (99.1% identity). Seven Maryland traps (24%) were positive for flaviviruses. These viruses were most similar to the *Calbertado* and *Nienokoue* flaviviruses, although the percent identity was very low (71.1%-74.3%) and they clearly separated from those viruses in phylogenetic analysis (**Figure 1 and Supplemental Figure 2**). These sequences likely derive from viruses that have not been sequenced yet but, since that they cluster with other mosquito flaviviruses (**Figure 1 and Supplemental Figure 2**), it is likely that they represent mosquito-infecting viruses rather than new human pathogens.

**Figure 1.**
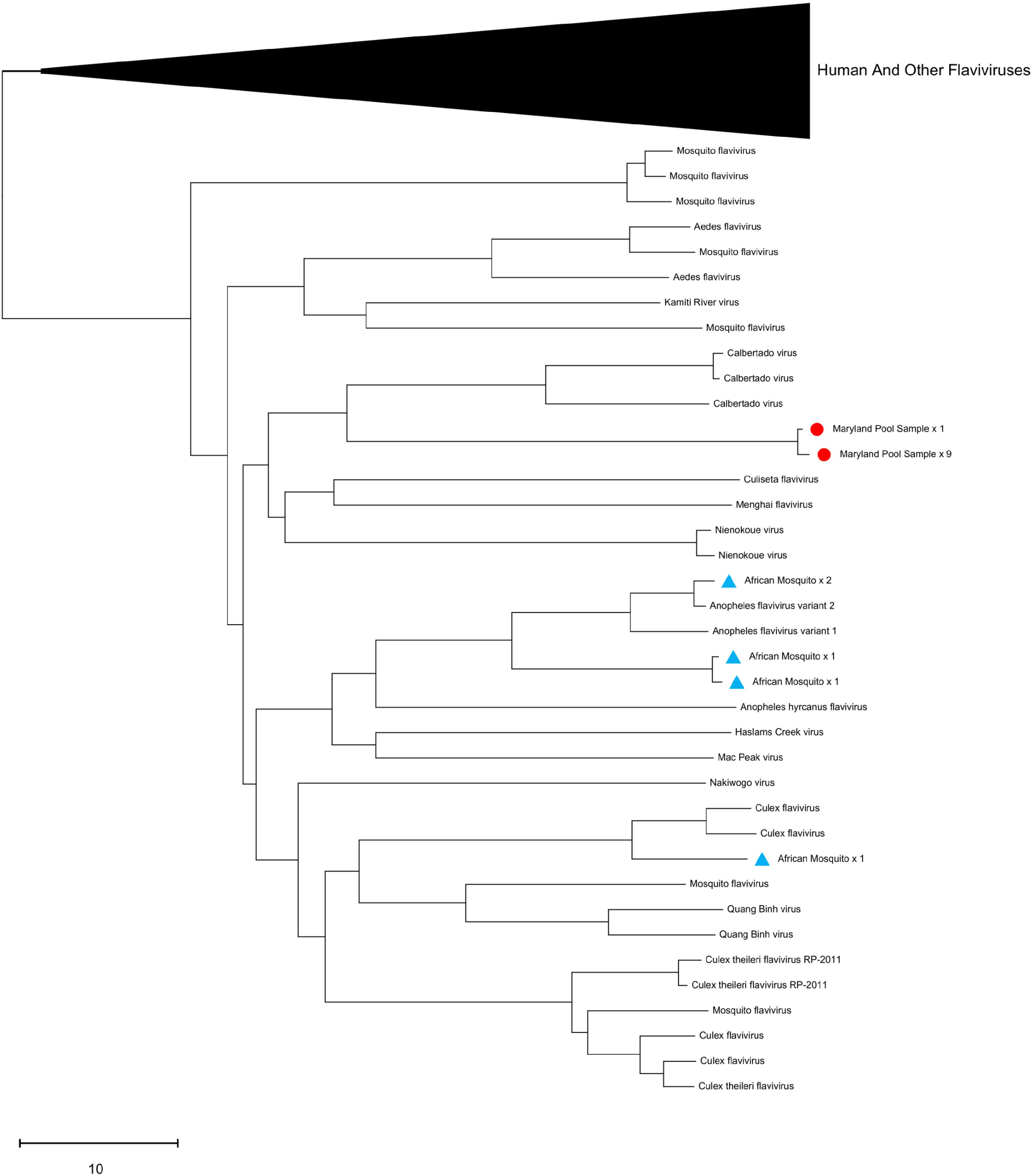
Phylogenetic analysis of flavivirus sequences amplified from mosquitoes. The neighbor-joining tree shows the relationships between the flavivirus sequences amplified from mosquito pools from Maryland (red circles) and from individual African mosquitoes (blue triangles). Phylogenetic tree without compressed branch available in **Supplemental Figure 2**.

One limitation of our study is that the mosquitoes collected in Maryland, USA were, as typical in many entomological surveys, stored at room temperature upon collection which might have affected RNA preservation. To assess the stability of viral and mosquito RNA in samples stored at room temperature, we kept pools of colony mosquitoes known to be infected with *Culex flavivirus* at room temperature for up to four weeks after collection, with and without preservative (ethanol or RNAlater). After RNA extraction and cDNA synthesis, we determined the amount of mosquito and virus RNA amplifiable using real-time PCR (see Material and Methods for details). Without preservative, the mosquito RNA was largely degraded after two weeks (detectable in only one of three replicates) and undetectable after four weeks (**Supplemental Figure 3**). By comparison, under the same conditions, viral RNA was still detectable after four weeks (**Supplemental Figure 3**). As expected, when the mosquitoes were preserved in either ethanol or RNAlater, neither viral nor mosquito RNA showed major change in concentration over four weeks at room temperature.

### Follow-up phylogenetic studies

The taxon-specific primers used in the high-throughput sequencing assay were designed to amplify all members of the chosen group while avoiding off-target amplification and providing as much taxonomic information as possible. However, these criteria, combined with the requirement for short sequences (to be sequenceable on a massively parallel sequencer) sometimes limits their resolution.

Thus, the Apicomplexa primers amplified multiple *Theileria* sequences but did not distinguish among species. We therefore amplified a longer DNA sequence (900 bp) of the 18S rRNA locus from the *Theileria*-positive African and Cambodian mosquitoes and sequenced them using Sanger sequencing technology. Phylogenetic analysis of these longer sequences, together with known *Theileria* species sequences deposited in NCBI, showed that the parasites amplified from the Cambodian mosquitoes were closely related to *T. sinensis,* while those from African mosquitoes were most closely related to *T. velifera* and *T. mutans* (**Figure 2**).

**Figure 2.**
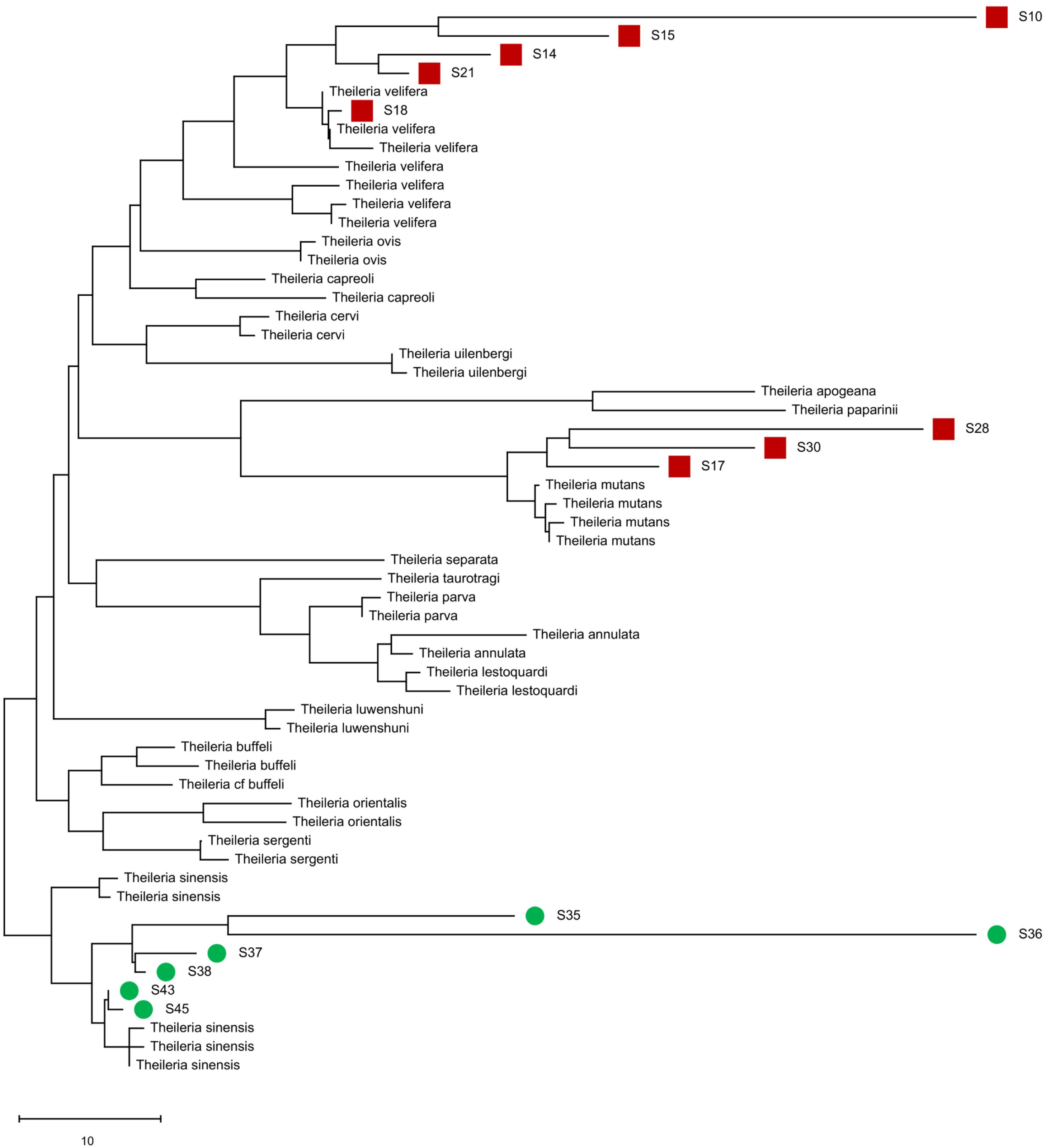
Phylogenetic analysis of *Theileria* sequences amplified from Cambodian and African *Anopheles* mosquitoes. The neighbor-joining tree shows the relationships between the 18S rRNA *Theileria* sequences amplified from samples positive by high-throughput sequences and those from known *Theileria* species deposited in NCBI. Sequences amplified from Cambodian mosquitoes are indicated in green circles, those amplified from African mosquitoes in red squares.

We also detected, in several African mosquitoes, filarial worm sequences whose taxonomic assignment was uncertain. One sequence was 100% identical to both *Loa loa* and *Dipetalonema sp. YQ-2006* (also known as *Mansonella*) while the sequence obtained from the same mosquitoes using a different primer pair was also most similar to *Dipetalonema* (*Mansonella*) but with 99.2% identity. To clarify the taxonomy of these sequences, we used PacBio long read technology to sequence 3.5kb of filarial worm mitochondrial DNA (amplified by long range PCR from these two mosquitoes). We compared these sequences to known filarial worm mitochondrial DNA sequences and found that these were most similar to, but distinct from, *Mansonella perstans* (94 and 96 nucleotide differences or^~^97.0% identity), while *Loa loa* was much more distantly related (^~^83.3% identity) (**Figure 3**). The genetic distance between *Mansonella perstans* and *Loa loa* in this tree was much higher (519 nucleotide differences, 83.5% identity) than using short amplicon data where these two sequences were identical, providing greater confidence in the phylogenetic analysis. We concluded that the filarial worms are most likely either *Mansonella perstans* or a very closely related species.

**Figure 3.**
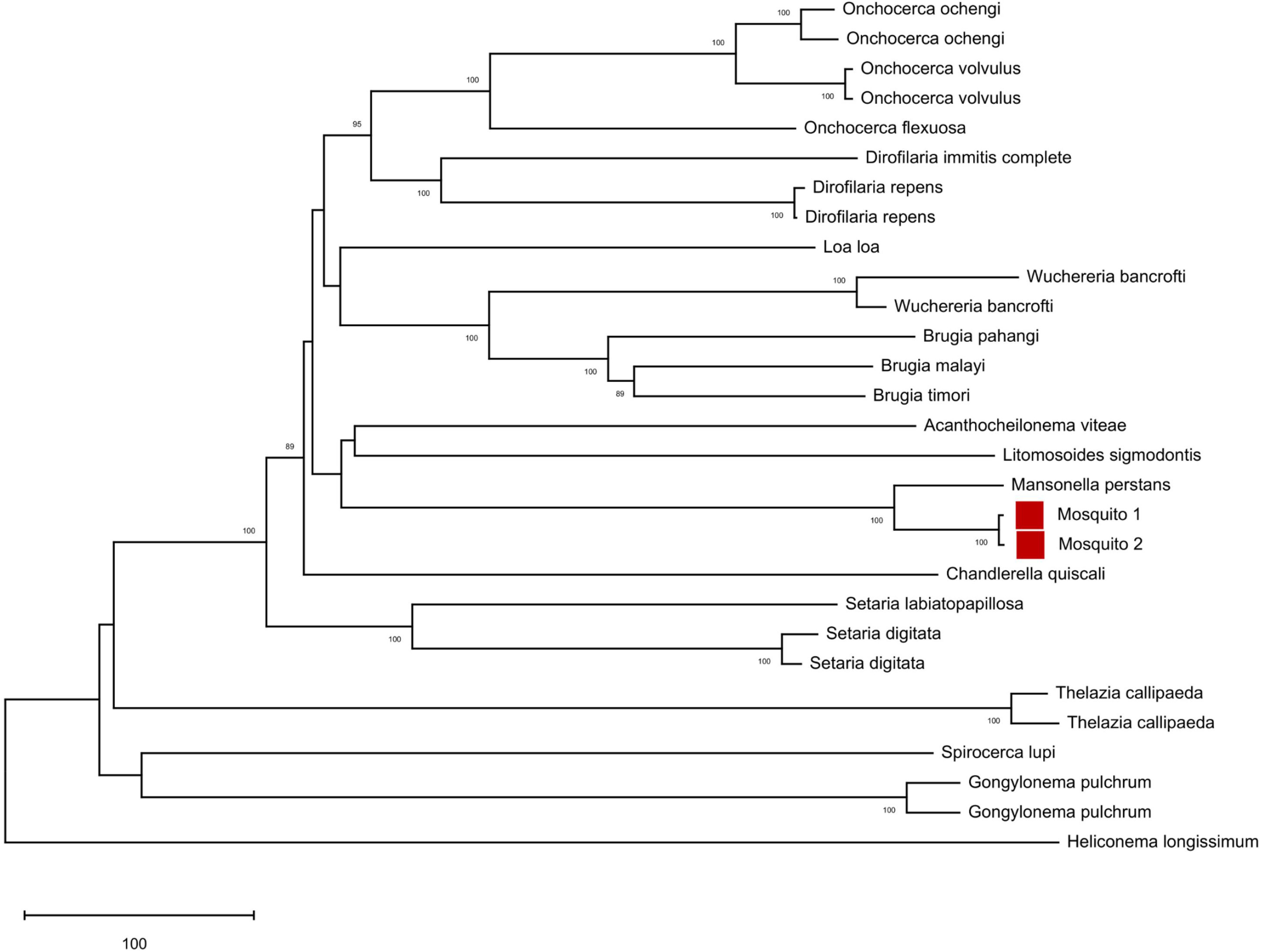
Phylogenetic analysis of unknown filarial worm sequences amplified from Guinean mosquitoes. The neighbor-joining tree shows the relationships between annotated filarial worm sequences and a 3.5 kb sequence amplified from two African mosquitoes (red squares) positive for filarial worms and sequenced using PacBio chemistry.

### Analysis of individual vs pooled mosquitoes

We analyzed both individual mosquitoes and pools of 50-291 mosquitoes. For the pools, 23 out of the 25 produced sequences demonstrating that the amplification of pools of up to 291 mosquitoes is feasible without significant PCR inhibition. Out of the 930 individual mosquitoes, 162 (17.4%) had at least ten reads from one or more parasites or arboviruses. By comparison, 23 out of the 25 traps (92%) yielded such sequences, meaning that fewer samples needed to be screened to detect pathogens when samples were pooled. Note however that the individual mosquitoes and the CDC traps were not collected from the same geographic locations and this could possibly cofound the results described.

### Analysis of DNA vs. RNA extracted from the same mosquito traps

All primers used for detecting parasites and viruses are located within single-exon genes and can amplify either DNA or cDNA with the same efficiency. However, the PCRs target genes that are typically highly expressed (e.g., ribosomal RNA genes) and we would therefore expect many more copies of RNA than DNA per cell, (although this could be diminished by the faster degradation of RNA molecules compared to DNA). To evaluate the relative sensitivity of our assay for screening DNA and RNA, we compared the results obtained by analyzing matched DNA and RNA isolated from the same mosquito pools. We found that for Spirurida, Kinetoplast, Microsporidia and *Plasmodium* PCR assays, 62.2% of the sequences identified were detected only in the cDNA sample and not in the corresponding DNA sample from the same trap. For those cases where a sequence was detected in both cDNA and DNA from a given trap, the cDNA yielded more reads in 89 of 119 instances (with, on average, 24.5-fold more reads). Read counts were higher in the DNA for only 29 of 119 cases with an average fold difference of 2.8 (one had equal read counts in cDNA and DNA). Out of these 29 cases, 22 (75.9%) of the sequences were most similar to *Trypanosoma* species despite these sequences representing only 17.4 of all sequences. On average, the cDNA samples produced 258 – 1,169 more reads per hit than the matching DNA samples (**Figure 3**), despite the storage of the samples at room temperature without preservative for more than 24 hours.

## Discussion

Vector-borne disease surveillance is an essential component of infectious disease control as it can enable rapid detection of outbreaks and guide targeted elimination efforts (e.g., through insecticide spraying). However, current approaches are extremely demanding in regards to human and financial resources, both for the sample collection and the identification of potential pathogens. Consequently, public health officials and vector biologists often have to focus on a handful of parasites associated with the most current threats. Current detection approaches also often lead to duplicated efforts, as different agencies interested in specific pathogens perform sample collection independently and have a high risk of failing to detect emerging pathogens until they cause outbreaks. Here, we describe application of a genomic assay that allows identification of a wide range of pathogens that can cause human and animal diseases, as well as of parasites of the vector that could potentially be useful as biological controls.

The analyses of several hundred mosquitoes collected in Cambodia, Mali, Guinea and Maryland revealed well-known human pathogens including *P. falciparum*, which was the target of the initial study of the Cambodian samples (24). In addition, we detected *Theileria* species and *Setaria digitata*, which cause livestock diseases in Southeast Asia (38–42). While we were initially unable to conclusively determine the exact *Theileria* species with our initial assay, targeted follow-up studies using longer amplicons and Sanger sequencing (**Figure 2**) revealed that the sequences amplified from the African mosquitoes were most closely related to *T. velifera* and *T. mutans,* which are both known to infect African cattle (43), whereas the Cambodian mosquitoes carried sequences most closely related to *T. sinensis,* a species that infects cattle in China (44).

*Theileria* parasites are transmitted by ticks, not mosquitoes, and the DNA sequences recovered likely derive from parasites taken up by the mosquitoes during a blood meal but likely not transmissible to another hosts. The *Schistosoma* species detected in mosquitoes from Africa also likely result from parasites present in a bloodmeal. In this regard, it is interesting to note that when one considers the samples collected in Maryland and analyzed with both DNA and RNA, the read counts (a proxy for the abundance of extracted molecules) for transmissible parasites (e.g., *Plasmodium*) or parasites of the mosquitoes (e.g., *Crithidia*, *Strigomonas* and *Takaokaspora*) were typically higher in the RNA samples than in the matched DNA samples while the opposite was true for parasites “sampled” during the blood meal but unlikely to develop in *Anopheles* mosquitoes (e.g., *Theileria*, *Trypanosoma*) (**Figure 4, Supplemental Figure 4 and Supplemental Table 6**). We speculate that this difference is due to the difference between developing, live, parasites still synthesizing RNA molecules and dead (possibly digested) parasites for which the RNA is slowly being degraded. Comparison of DNA and RNA from the same mosquito could perhaps provide a tool to differentiate transmissible parasites from those sampled by the vector.

**Figure 4.**
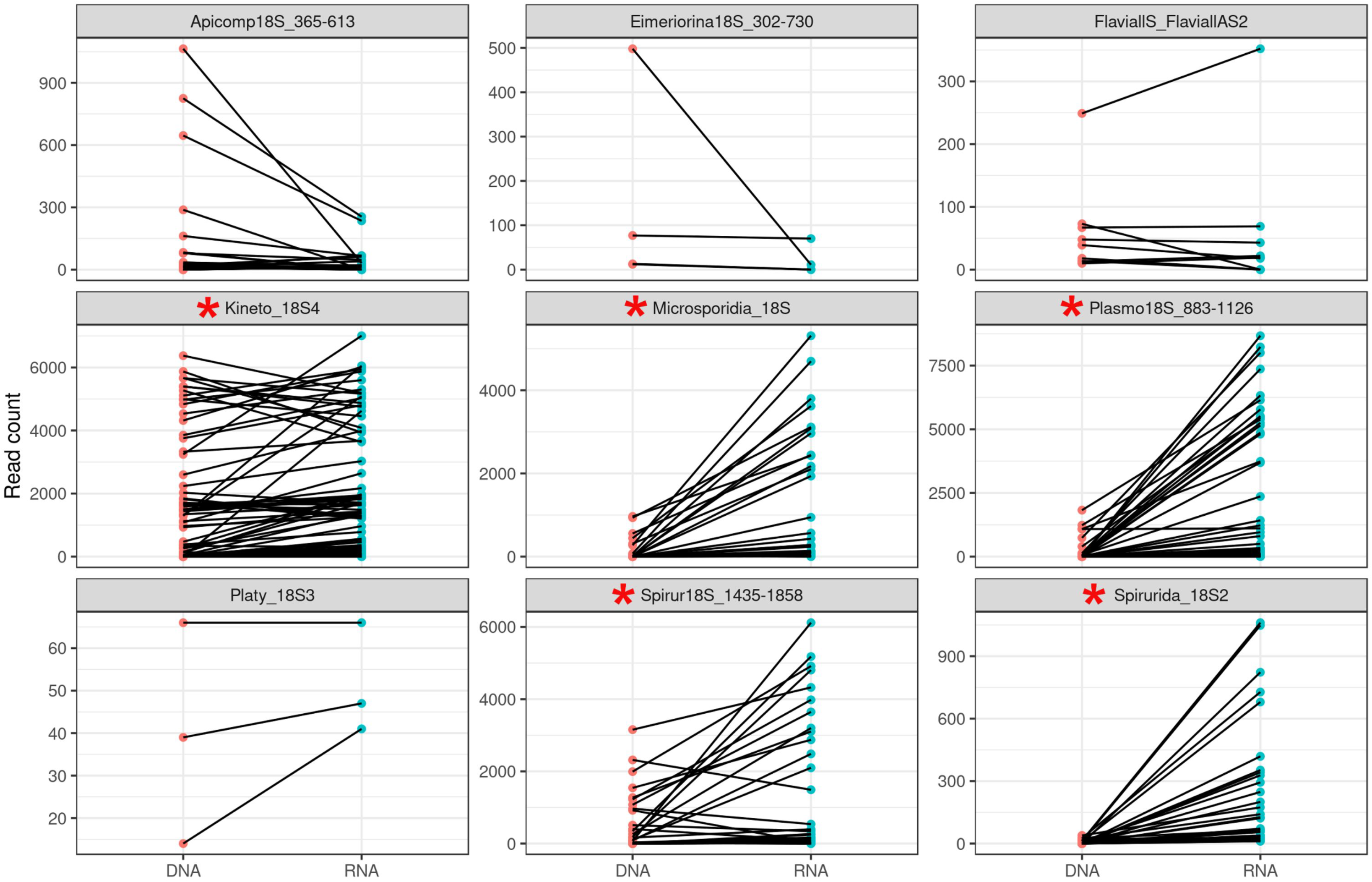
Comparison of the number of reads obtained for different taxa from matched DNA and cDNA samples derived from Maryland mosquito pools. Each panel represents results from one primer set and each pair of points connected by a line shows the number of reads matching a single species detected in both the DNA (left) and RNA (right) from the same sample. For five primers (red asterisks), the RNA samples yield significantly more reads than the matching DNA samples (p < 0.05, Bonferroni-corrected pairwise t-tests).

We also identified, in two African mosquitoes, sequences similar to known filarial worms but identical to multiple sequences present in the database. Using this information, we characterized longer DNA sequences and showed that these two mosquitoes likely carried *Mansonella perstans* parasites. Since the PCR primers are designed to amplify any member of the selected taxa, they can reveal the presence of novel pathogens as long as they are phylogenetically related to known parasites. This feature is a key advantage of our assay for vector-borne disease surveillance as it may enable early detection of emerging pathogens and zoonoses and provide a basis for rapid response.

In addition to known human parasites and potential emerging pathogens, this single-stop assay also provides another source of information valuable for vector-borne disease control: 9% of the individual mosquitoes and 62% of pooled mosquito samples screened yielded sequences of microsporidians related to well-characterized arthropod parasites, which could potentially be used to guide the development of targeted biological vector control. This ability to detect multiple parasites at once in a high-throughput manner and across a wide range of taxonomical groups could reduce duplication of collection efforts and costs, as mosquitoes collected for one purpose could be screened for many parasites affecting both humans and animals. In addition, comprehensive characterization of the parasites present in a given mosquito may also improve our understanding of the general factors regulating infection and transmission: several studies have shown that immunity and previous infections can influence the response of mosquitoes to human parasites and their transmission (45–47) and information of current infections of wild-caught mosquitoes could, for example, significantly improve our assessment of their vector capacity.

Several of the infectious diseases that have recently caused major public health challenges by spreading outside of their typical range(7, 8) or emerging as novel human infectious diseases(5, 6), are caused by viruses transmitted by mosquitoes. We therefore extended our assay to capture, using the same approach as for eukaryotic parasites, both known and novel flaviviruses. Since flaviviruses are RNA viruses and RNA degrades much faster than DNA, we first examined how nucleic acid degradation influenced our ability to detect virus over time. To test RNA preservation, we collected mosquitoes known to carry *Culex flavivirus* and isolated RNA from pools of five mosquitoes, either immediately frozen or kept at room temperature for two or four weeks, with either no preservative, ethanol or RNAlater. The mosquitoes stored in preservatives had minimal loss of viral (and mosquito) RNAs as determined by qRT-PCR (**Supplemental Figure 3**). Even when stored without preservatives, viral RNA were detectable after 4 weeks at room temperature (although with a reduction of, on average, 10.7 PCR cycles), demonstrating a remarkable stability of the RNA, possibly due to protection provided by the viral capsid (by contrast very little mosquito RNA remained amplifiable after two weeks at room temperature, **Supplemental Figure 3**). As a proof-of-principle and to demonstrate the potential of this approach for viral disease surveillance, we screened the Maryland mosquito pools and the individual African mosquitoes for flaviviruses. We identified several viruses, distinct from known viruses (**Figure 1**) and, based on their phylogenic position, likely to infect mosquitoes rather than humans.

Based on the results described above, we believe that this single high-throughput assay can provide a wide range of information critical for vector-borne disease researchers and public health officials. However, several limitations need to be noted. One caveat is that, whereas false positive detection of a species is highly unlikely (aside from laboratory cross-contamination), several factors could lead to false negatives. Thus, while the primers were designed to amplify all known sequences of a given taxon as effectively as possible, nucleotide differences at the primer binding sites could prevent efficient amplification of a specific species. This potential problem could be particularly problematic if several related parasites are present in the same sample but are differentially amplified: for example, it could be possible that a *Plasmodium* parasite might be mis-detected if the sequences generated by an Apicomplexan primer pair are out competed by *Theileria* sequences. Similarly, poor preservation of the nucleic acids in one sample could also lead to false negatives. False negatives could also occur for stochastic reasons: if only a few parasite cells are present in one sample (*e.g.*, an *Anopheles* mosquito infected by a *Plasmodium* ookinete) it is possible that no DNA will be present in the PCR reaction (especially if the extract gets divided across many reactions). One approach to circumvent this limitation could be to test cDNA instead of DNA (48, 49): our analyses of the Maryland mosquitoes showed that, for many primer sets, amplification of cDNA resulted in higher read counts than amplification of DNA extracted from the same samples, despite the sub-optimal preservation of these samples. Another limitation is the specificity of taxonomic assignment. As discussed above, if the sequenced amplicon does not contain enough information to distinguish similar species, subsequent experiments may be required to confirm pathogen identity for important detection events.

Finally, we showed that analyses of fairly large pools of mosquitoes (up to ^~^300 mosquitoes) were possible with our assay. This feature could be extremely useful in specific situations, such as for efficiently detecting emerging pathogens, monitoring the spread of pathogens into new regions, or for validating the success of elimination control programs.

## Conclusion

This study demonstrates how our high-throughput, one-stop assay could efficiently complement current toolkits to prevent vector-borne diseases by providing a comprehensive description of known and emerging human viruses and parasites, informing on animal pathogens that could affect a region’s economy, and indicating possible biological control candidates that could be used against these disease vectors. One additional feature of this sequencing-based assay is the ease of customizing it to different settings and research questions. Since the assay relies on PCR primers, it is straightforward to add and remove primers for specific taxa of interest, or to combine them with additional PCRs to characterize, for example, the source of the blood meal (27).

## Abbreviations

CNM: National Center for Parasitology, Entomology, and Malaria Control
ATCC: American Type Culture Collection

## Acknowledgements

We wish to thank our collaborators at the National Center for Parasitology, Entomology, and Malaria Control (CNM) in Phnom Penh, Stop Palu and the National Malaria Control Program in Guinea and at the Malaria Research and Training Center, University of Science, Techniques and Technologies, Bamako, Mali for their help with the sample collection and processing, as well as Erin Haser, Rachel Koch, Jessica Moskowitz, Rachel Shuck, and Elizabeth Allan. We also thank Stop Palu (RTI), the U.S. President’s Malaria Initiative, and all those participating in collections for support in the sample collection and processing in Guinea. The findings and conclusions in this report are those of the authors and do not necessarily represent the official position of the Centers for Disease Control and Prevention.

## Availability of data and material

The datasets generated for the current study are available in the NCBI SRA repository (Accession numbers SRR12797126 - SRR12797220, SRR12797360 - SRR12797683, SRR12796164 - SRR12796923 and SAMN16182375 - SAMN16183134)

## Funding

This study was supported by DS and by the Intramural Research Program (NIAID-NIH) to BSTL. SRI was funded by the U.S. President’s Malaria Initiative.

## Notes

**Conflict of interest statement:** The authors have stated explicitly that there are no conflicts of interest in connection with this article.

